# Phenome-wide association studies (PheWAS) across large “real-world data” population cohorts support drug target validation

**DOI:** 10.1101/218875

**Authors:** Dorothée Diogo, Chao Tian, Christopher S. Franklin, Mervi Alanne-Kinnunen, Michael March, Chris C. A. Spencer, Ciara Vangjeli, Michael E. Weale, Hannele Mattsson, Elina Kilpeläinen, Patrick M.A. Sleiman, Dermot F. Reilly, Joshua McElwee, Joseph C. Maranville, Arnaub K Chatterjee, Aman Bhandari, the 23andMe Research Team, Mary-Pat Reeve, Janna Hutz, Nan Bing, Sally John, Daniel MacArthur, Veikko Salomaa, Samuli Ripatti, Hakon Hakonarson, Mark J. Daly, Aarno Palotie, David Hinds, Peter Donnelly, Caroline S. Fox, Aaron Day-Williams, Robert M. Plenge, Heiko Runz

**Affiliations:** Merck Sharp & Dohme, Kenilworth, NJ, USA.; 23andMe Inc., Mountain View, CA, USA.; Genomics Plc, Oxford, UK.; Institute for Molecular Medicine Finland (FIMM), University of Helsinki, Helsinki, Finland.; The Children’s hospital of Philadelphia and University of Pennsylvania, Philadelphia, PA, USA.; National Institute for Health and Welfare, Helsinki, Finland.; Eisai, Andover, MA, USA.; Pfizer, Cambridge, MA, USA.; Biogen, Research and Early Development, Cambridge, MA, USA.; Broad Institute of MIT and Harvard, Cambridge, MA, USA.; Analytic and Translational Genetics Unit, Department of Medicine, Massachusetts General Hospital, Boston, MA, USA.; Psychiatric & Neurodevelopmental Genetics Unit, Department of Psychiatry, Massachusetts General Hospital, Boston, MA, USA.; Department of Neurology, Massachusetts General Hospital, Boston, MA, USA.

## Abstract

Phenome-wide association studies (PheWAS), which assess whether a genetic variant is associated with multiple phenotypes across a phenotypic spectrum, have been proposed as a possible aid to drug development through elucidating mechanisms of action, identifying alternative indications, or predicting adverse drug events (ADEs). Here, we evaluate whether PheWAS can inform target validation during drug development. We selected 25 single nucleotide polymorphisms (SNPs) linked through genome-wide association studies (GWAS) to 19 candidate drug targets for common disease therapeutic indications. We independently interrogated these SNPs through PheWAS in four large “real-world data” cohorts (23andMe, UK Biobank, FINRISK, CHOP) for association with a total of 1,892 binary endpoints. We then conducted meta-analyses for 145 harmonized disease endpoints in up to 697,815 individuals and joined results with summary statistics from 57 published GWAS. Our analyses replicate 70% of known GWAS associations and identify 10 novel associations with study-wide significance after multiple test correction (P<1.8x10^-6^; out of 72 novel associations with FDR<0.1). By leveraging directionality and point estimate of the effect sizes, we describe new associations that may predict ADEs, e.g., acne, high cholesterol, gout and gallstones for rs738409 (p.I148M) in *PNPLA3*; or asthma for rs1990760 (p.T946A) in *IFIH1*. We further propose how quantitative estimates of genetic safety/efficacy profiles can be used to help prioritize candidate targets for a specific indication. Our results demonstrate PheWAS as a powerful addition to the toolkit for drug discovery.

**One Sentence Summary:** Matching genetics with phenotypes in 800,000 individuals predicts efficacy and on-target safety of future drugs.

## INTRODUCTION

The discovery and development of novel therapeutics is difficult. It may take 15 years to advance a new molecular entity from therapeutic hypothesis to approval, with development costs in the billion dollar range and only 10% chance of a new drug tested in humans eventually getting approval^1^. Two reasons stand out to explain the high failure rate of clinical trials and receding return on R&D investment across the pharmaceutical industry: A lower efficacy of the compound in the targeted disease population than anticipated from preclinical studies; and the occurrence of unintended drug effects, particularly mechanism-based adverse drug events (ADEs) uncovered only in late-stage clinical trials^2^. A greater understanding of human data relevant to the drug target at early stages of drug development is generally considered to increase the probability of success^1, 3, 4^.

Resources that systematically capture biomedical information on vast numbers of individuals are revolutionizing our ability to understand the complexities of human biology and morbidity. Electronic health records (EHRs) and similar “Real-World Data” (RWD) have rapidly become well-established tools for epidemiological and post-marketing research^5, 6^. Recently, a surge of initiatives have sought to link such phenotype resources with genome-scale genetic data in order to gain further insights into the genetics of common diseases^7, 8, 9, 10, 11, 12, 13, 14^.

An attractive approach for such genotype-phenotype resources to help accelerate drug development is through Phenome-Wide Association Studies (PheWAS). PheWAS are an unbiased approach to test for associations between a specific genetic variant and a wide range of phenotypes in large numbers of individuals^7, 15^. By exploring the associations of a genetic variant that impacts the function of a drug target gene, PheWAS in RWD cohorts may enrich the drug discovery process for five reasons: 1) association studies in RWD cohorts may validate target-disease links in cohorts that more closely resemble the “real-world”, i.e. the patients that will ultimately receive a drug^16^; 2) by unraveling pleiotropy, PheWAS may improve our understanding of the biological functions of a target, or hint at concealed pathophysiological connections between disease entities previously considered as distinct^17, 18^; 3) PheWAS may reveal opportunities for drug repurposing, an attractive alternative to *de novo* drug development^19, 20^; 4) PheWAS may point to phenotypes that associate with an inverse directionality of target function, thus unravelling potential ADEs at very early stages of a developmental program, minimize risks to trial participants, and help define the most appropriate patient populations to benefit from a drug^20^; 5) through quantitative estimates from genetic safety and efficacy profiles, PheWAS may help prioritize multiple possible targets by identifying the target with the most promising therapeutic window. Despite these benefits the ability for PheWAS to substantially add to the decision making in drug development is thwarted by the difficulty to obtain and systematize comprehensive genotypes and phenotypes across very large numbers of individuals.

Here, we have tested the hypothesis that PheWAS can inform target validation at early stages of drug development. We selected candidate drug targets across a range of therapeutic indications based on their support from genome-wide association studies (GWAS). To maximize power, we mapped a large spectrum of clinical endpoints from four of the world’s largest RWD population cohorts and conducted association testing in up to 697,815 individuals. Our results show that PheWAS, despite limitations, enrich drug discovery with valuable information.

## RESULTS

### Assessing pleiotropy for SNPs in/near 19 drug targets through meta-analyses across four Real-World Data cohorts

In this study, we queried the literature for genes nominated through GWAS as putatively causally linked to the risk for common complex human diseases and supported by various degrees of additional genetic or biological evidence. We selected 19 genes that, based on previously described genetic associations with either immune-mediated (9 genes: *ATG16L1, CARD9, CD226, CDHR3, GPR35, GPR65, IFIH1, IRF5, TYK2*), cardiometabolic (8 genes: *F11, F12, GDF15, GUCY1A3, KNG1, LGALS3, PNPLA3, SLC30A8*) or neurodegenerative diseases (2 genes: *LRRK2, TMEM175*), were evaluated as potential novel drug targets. Gene-disease associations had been established through 25 common lead single nucleotide polymorphisms (SNPs) that all reached a conservative level of statistical significance (P<5x10^-8^) for association in GWAS with at least one phenotype of relevance to drug discovery and development (**Table S1**). All of these SNPs have either been demonstrated to impact the target gene in functional studies (genetic evidence), or locate proximal to a gene implicated in a biological mechanism related to the GWAS phenotype (biological evidence). Our selection ranged from targets with little biological knowledge beyond GWAS nomination (e.g. *TMEM175* for Parkinson’s disease) to targets with drug candidates in early clinical trials (e.g. *F11* for thromboembolism). Details on the genetic and biological support for all selected genes and SNPs is provided in **Supplementary Information**.

To broadly investigate pleiotropic effects of the 25 chosen SNPs in a maximal number of individuals, we interrogated four large RWD cohorts that link genome-wide genotype data from individuals of European ancestry with extensive phenotypic data: the 23andMe Inc. cohort with self-reported phenotypes on 671,151 research participants^21^, the interim UK Biobank cohort analyzed by Genomics plc with questionnaire-based health information on 112,337 participants (from the first genetic data release in May 2015)^10^, and two EHR-based cohorts from an adult Finnish cohort (FINRISK; 21,371 participants)^22^ and from a pediatric healthcare population from the Children’s Hospital of Philadelphia (CHOP; 12,044 patients)^23^ (**Table 1** and Methods). All four cohorts contributed phenotypic data in different formats (medical interviews, self-reports, WHO ICD codes, or ICD9-CM codes) in both shared and distinct phenotype categories (**Fig. 1A**). Together, the four RWD cohorts allowed association testing for a total of 1,892 binary endpoints. Manual phenotype mapping identified 145 distinct clinical endpoints that could be reliably harmonized across two or more cohorts, enabling meta-analyses in up to 697,815 individuals (**Fig. 1B** and **Table S2**). As illustrated in **Fig. 1C**, these 145 mapped phenotypes represent a broad spectrum of disease categories and, as typically observed in RWD, show significant variability in the case:control ratios, both within and between cohorts.

**Fig. 1.**
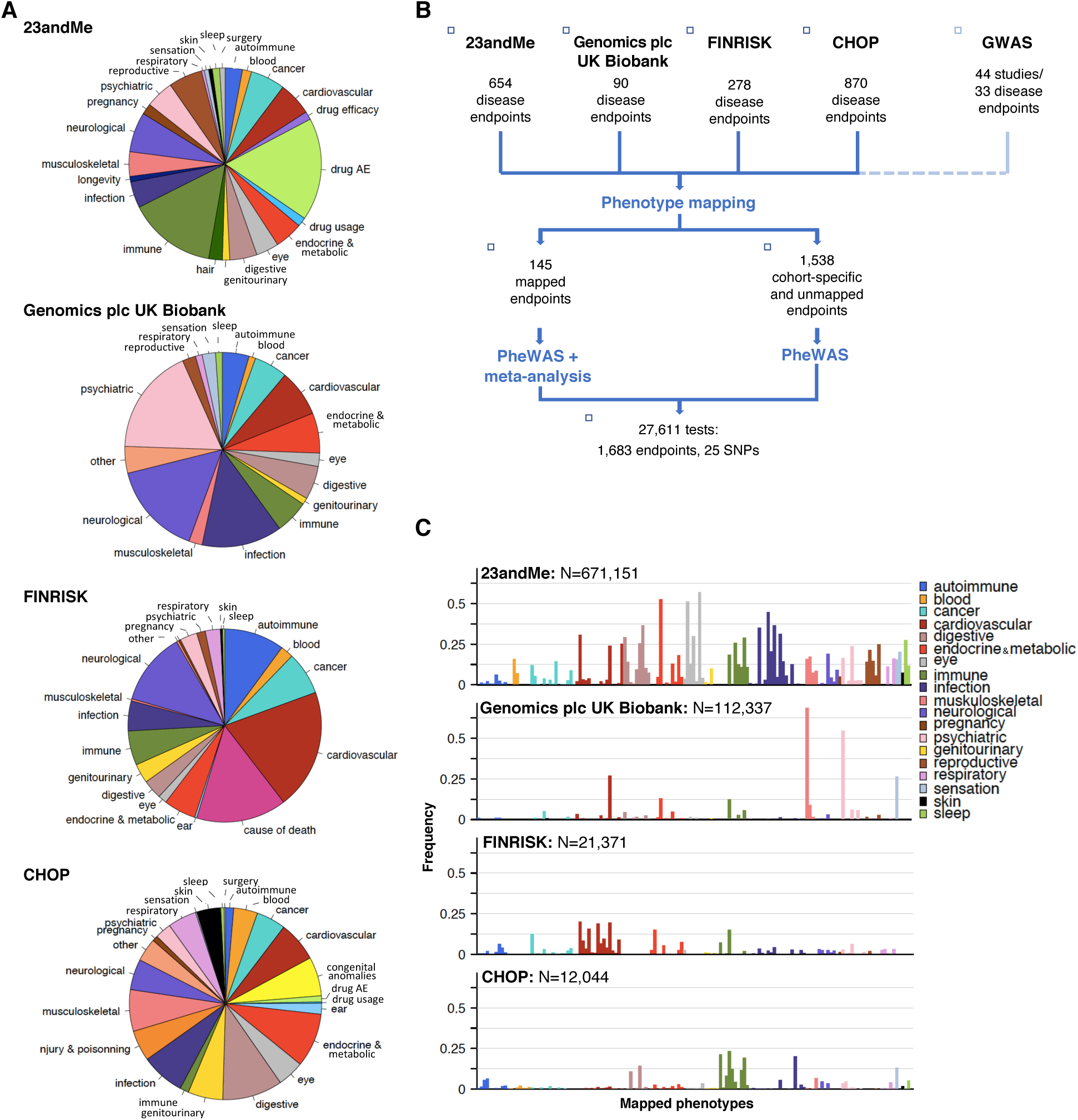
Phenotypes tested and study design. (A) Categories of phenotypes assessed in the 23andMe, Genomics plc UK Biobank, FINRISK and CHOP RWD cohorts. **(B)** Manual phenotype mapping was performed to identify phenotypes shared between cohorts. One hundred and forty-five phenotypes were captured with at least 20 cases in at least 2 cohorts. After PheWAS in each cohort separately, the 145 phenotypes were meta-analyzed to increase statistical power and enable systematic comparisons of results between cohorts. **(C)** The 145 mapped phenotypes (see **Supplementary Table 2**) represent a broad spectrum of phenotypic categories and are captured with variable case:control ratios in the cohorts tested.

**Table 1.** Cohorts included in this study.

| **Cohort** | **Participants geographic distribution** | **Phenotypes source** | **N binary endpoints tested \*** | **Max sample size** |
| --- | --- | --- | --- | --- |
| 23andMe | 89% USA (adult) | Questionnaire-based self-reports | 654 | 671,151 |
| Genomics plc UK Biobank | 100% UK (adult) | Questionnaire-based self-reports, medical interviews and follow-up | 90 | 112,337 |
| FINRISK | 100% Finns (adult) | National health registries (ICD8,9,10) | 278 | 21,371 |
| CHOP | 100% USA (pediatric) | Electronic health records (ICD9-CM) | 870 | 12,044 |
| Genomics plc GWAS | mixed | Mixed – multiple independent disease-specific cohorts | 34 | - |

*\* Number of binary endpoints with $N$ cases $\geq 20$*

### Association testing in RWD cohorts validates known GWAS signals

We first evaluated whether association testing in the four RWD cohorts (referred to as ‘RWD PheWAS’) replicated established results from published GWAS. GWAS had associated the 25 tested SNPs with genome-wide significance to 58 binary disease endpoints. Of these, 30 endpoints were ascertained with adequate power (beta ≥0.8) to reach FDR<0.1 (P<3.8x10^-4^) in the RWD cohorts. We observed that 21 of the 30 (70%) powered GWAS associations replicated (FDR<0.1) in our RWD PheWAS meta-analysis (**Fig. S1** and **Table S3**). As expected from data obtained in real-world settings, the replication rate of known associations was highly disease-dependent. For instance, out of the 9 associations that failed to replicate despite sufficient case numbers in the cohorts, 6 were associations with inflammatory bowel disease (IBD), Crohn’s disease (CD) or ulcerative colitis (UC), likely reflecting suboptimal ascertainment of these endpoints in real-world settings. Nonetheless, the high replication rate of previously reported associations demonstrates the power of combining disease-agnostic RWD cohorts from various sources to detect and validate true SNP-disease associations, and to substantiate therapeutic hypotheses.

### Meta-PheWAS across RWD cohorts identify novel SNP-endpoint associations

We next investigated whether meta-PheWAS across the four RWD cohorts could identify novel associations to support the proposed clinical indication, suggest alternative indications for drug repositioning, or uncover potential target-related ADEs. To improve statistical power in this analysis, the RWD meta-PheWAS results were further combined with summary statistics from published GWAS studies of 34 diseases available from a larger database assembled and harmonized by Genomics plc (referred to as Genomics plc GWAS, **Supplementary Information**). Overall, 27,763 association tests (across 145 harmonized and 1,538 cohort-specific endpoints) resulted in 10 putative novel associations reaching study-wide significance after Bonferroni correction (P<1.8x10^-6^) (**Table 2**). Using a less stringent significance threshold of FDR<0.1 (P<7x10^-4^) previously applied in PheWAS^24^, we identified 72 distinct putative novel associations (**Fig. 2**, **Fig. S2**, **Table S4** and **Supplementary Datasheet)**. Forty-four of these putative novel associations showed directions of effect consistent with the proposed clinical indication for a drug and may hint at potential repositioning opportunities. Conversely, 27 showed directions of effect opposite to the proposed clinical indication and may suggest safety signals that could endanger therapeutic success and warrant monitoring for in preclinical models and clinical trials.

**Fig. 2.**
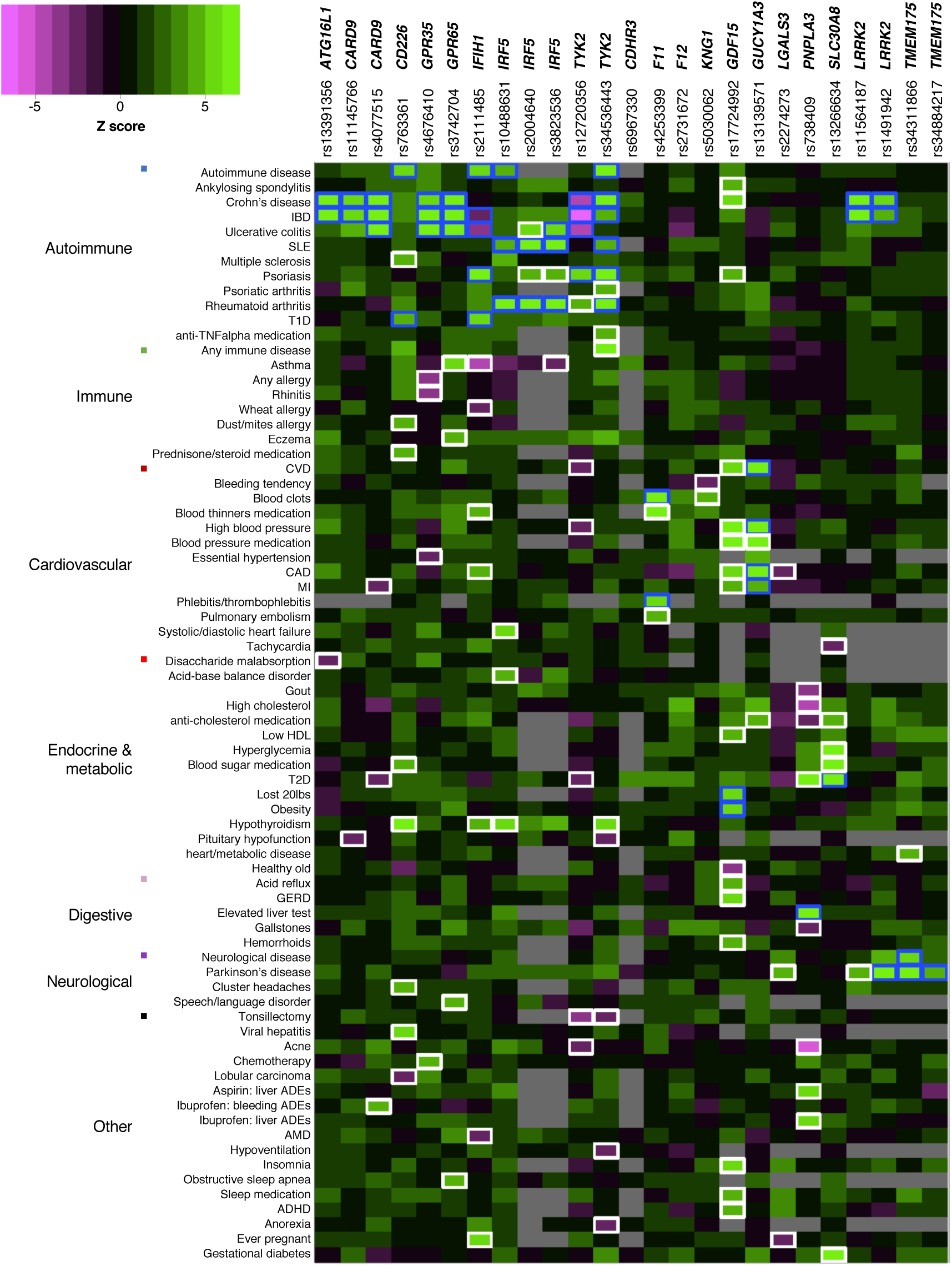
PheWAS results for 25 GWAS SNPs in/near candidate drug targets from meta-analysis of 4 RWD cohorts with published GWAS data. Phenotypes associated at FDR<0.1 (P<7e-4) with at least one SNP in the meta-PheWAS are represented. Direction of effect of the known disease-risk increasing allele related to the therapeutic hypothesis is indicated. A positive Z-score (in green) indicates increased risk, a negative Z-score (in purple) indicates reduced risk. Known and novel associations reaching FDR<0.1 are outlined in blue and white respectively.

**Table 2.**
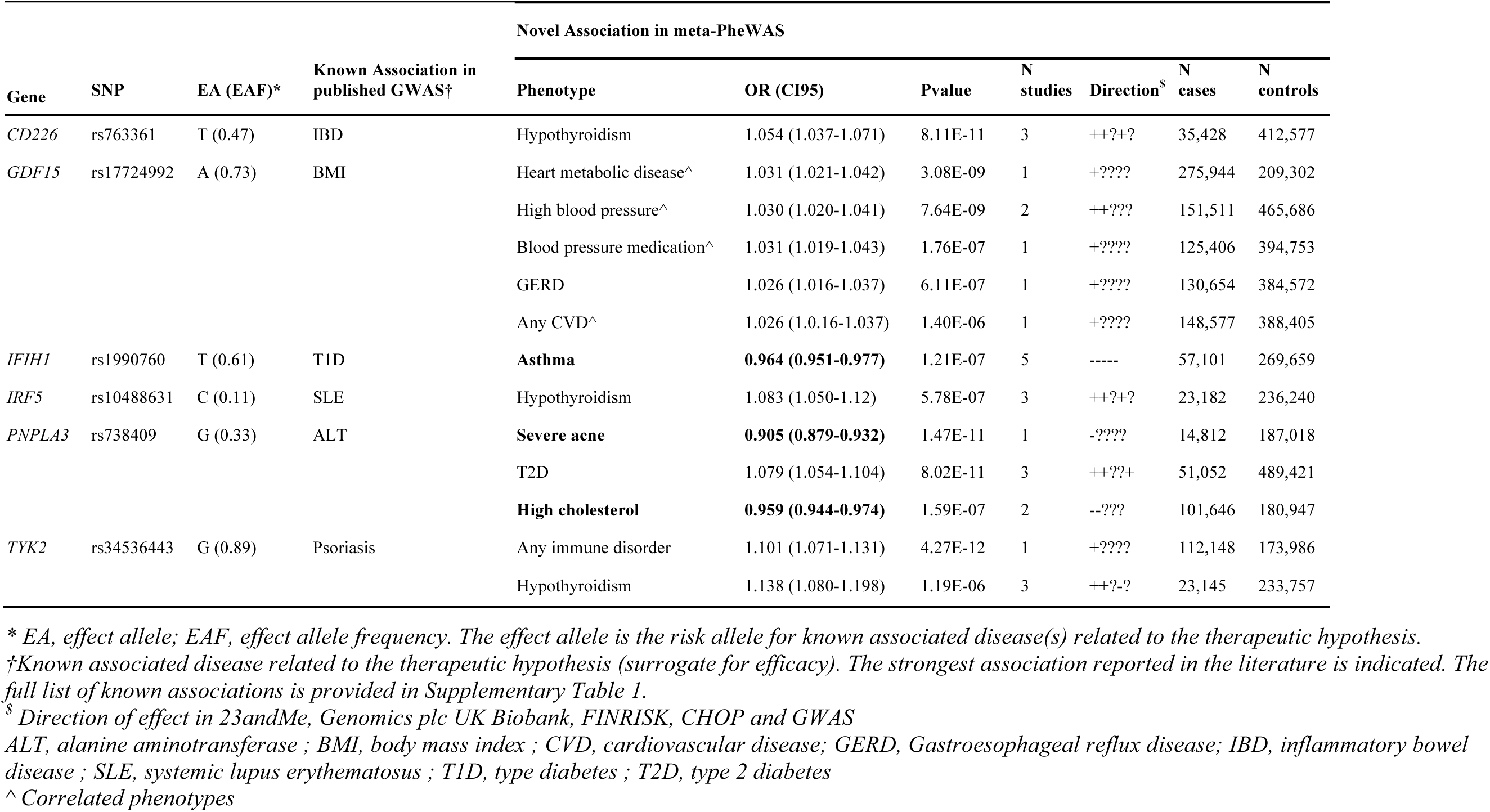
**Significant novel associations in the PheWAS meta-analysis.** Associations reaching P<1.8e-6 (Bonferroni–corrected significance threshold) in the meta-analysis of RWD PheWAS results with GWAS results are shown. The full list of potential novel SNP-phenotype pairs reaching FDR<0.1 is provided in **Table S4**. The effect of the allele increasing the risk for known associated disease(s) supporting the therapeutic hypothesis is shown. Novel associations with direction of effect opposite to the known associated disease(s) effect are highlighted in bold.

### Apparent pleiotropy and endpoint co-morbidities challenge target validation through PheWAS

A challenge to the PheWAS approach is to reliably distinguish true pleiotropic associations of a SNP (or SNPs in strong LD with the lead SNP), suggesting a shared causal mechanism, from unrelated associations driven by independent SNPs at a locus^17^. For instance, in our meta-PheWAS, the putative association of rs2274273 near *LGALS3* (encoding the galactin-3 protein) with Parkinson’s disease (PD) (OR=0.94, P=1x10^-4^) likely reflects a distinct causal mechanism previously attributed to *GCH1*^25^. rs2274273 is a protein quantitative trait locus (pQTL) that controls plasma levels of galectin-3^26^. Through a Bayesian test for co-localization using summary statistics from published GWAS studies^27, 28^, we excluded rs2274273 as a causal SNP for PD (posterior probability for a shared variant leading the PD and galectin-3 levels associations = 0.0008%) (**Fig. S3**).

A second challenge to PheWAS is the existence of common co-morbidities among endpoints, or alternatively an insufficient distinction between phenotypes^18^. In our meta-PheWAS, rs17724992 near *GDF15* showed association with multiple cardiovascular-related phenotypes, which is likely mediated by the known association of this SNP with body mass index (BMI)^29^, an established risk factor for cardiovascular disease^30^. This is supported by the lack of association of rs17724992 with coronary artery disease (CAD) in the large GWAS published by the CARDIoGRAMplusC4D consortium (P=0.17)^31^. Follow-up customized association analyses adjusting for specific phenotypic covariates are required to distinguish true pleiotropic effects and inform target validation.

In summary, these two examples demonstrate that thorough investigation of association results can reduce biases introduced through PheWAS.

### Pleiotropy of rs738409 (p.I148M) predicts a risk for multiple potential ADEs upon inhibiting *PNPLA3*

Among the 10 study-wide significant associations, our meta-PheWAS revealed multiple novel associations for the *PNPLA3* missense SNP rs738409 (p.I148M). The rs738409-G allele has previously been reported as associated with an increased risk for non-alcoholic fatty liver disease (NAFLD), alcohol-related cirrhosis and hepatic steatosis, as well as elevated alanine aminotransferase (ALT) levels, most likely through a gain-of-function (GOF) mechanism (**Supplementary Information**). Consistent with these findings, our meta-PheWAS found rs738409-G to be associated with elevated liver tests (OR=1.25, P=4x10^-45^) (**Fig. S4**). Beyond that, our analysis also indicated that carriers of the rs738409-G allele that increases ALT are more prone to develop liver toxicities when treated with nonsteroidal anti-inflammatory drugs (NSAIDs) such as ibuprofen (OR=1.43, P=4.6x10^-5^) or aspirin (OR= 1.57, P=5.3x10^-5^). In addition, the meta-PheWAS revealed significant associations between rs738409-G and an increased risk of T2D (OR=1.08, P=8x10^-11^), as well as a decreased risk for acne (OR=0.90, P=1.5x10^-11^), high cholesterol (OR=0.96, P=1.6x10^-7^) or the intake of cholesterol-lowering medications (OR=0.97, P=2x10^-4^), gout (OR=0.92, P=4.1x10^-5^), and gallstones (OR=0.95, P=2.7x10^-4^). All these associations remained prominent after adjusting for elevated liver tests (**Table S5**). Associations of rs738409-G with T2D and high cholesterol were supported by independent recent data from the GoT2D 82k exome chip study (OR=1.06, P=7.7x10^-5^) and the LDL GLGC 300K exome chip study (Beta =-0.018, P=1 x10^-8^), where fine-mapping confirmed rs738409 to be the most likely causal SNP^32, 33^. Taken together, our PheWAS results support the hypothesis that therapeutic inhibition of PNPLA3 could treat liver diseases. They also support T2D as a potential alternative indication for PNPLA3 inhibition. However, concomitant inverse associations with multiple other endpoints, including acne and high plasma cholesterol levels, indicate potential clinically relevant on-target ADEs that should be considered in decisions to progress PNPLA3 inhibitors towards clinical development.

### *IFIH1* partial loss-of-function increases the genetic risk for asthma

As another example of pleiotropic effects identified in our meta-PheWAS, carriers of the *IFIH1* (encoding MDA5) rs1990760-C allele (MAF=40%) have an established lower risk for several autoimmune diseases (type 1 diabetes, T1D; vitiligo; systemic lupus erythematosus, SLE; psoriasis) and an increased risk for ulcerative colitis (UC) (**Supplementary Information**). Functional studies suggest that rs1990760-C (p.T946A) causes *IFIH1* loss-of-function (LOF), and additional *IFIH1* LOF alleles have been shown to protect against T1D, vitiligo, psoriasis and psoriatic arthritis (PsA) (**Supplementary Information**). Our meta-PheWAS support these associations (**Fig. 2** and **Table S3**). Importantly, we also found a significant novel association between rs1990760-C and increased risk for asthma (OR=1.04, P_meta_=9.0x10^-8^) (**Fig. 3A**). The association between rs1990760 and asthma was supported by data from all four RWD cohorts as well as the GABRIEL and EVE asthma GWAS cohorts (OR_meta_=1.04, P_meta_=6.5x10^-8^)^34, 35^, despite lack of power to detect an association with rs1990760 in the published GWAS cohorts alone (**Fig. 3B**). This association remained significant after adjustment for autoimmune diseases in the 23andMe cohort, demonstrating that the asthma association is independent of the previously established associations of rs1990760 with autoimmunity (**Table S6**). Co-localization analysis confirmed that the same SNP was responsible for the SLE, UC and asthma associations at the locus, supporting true pleiotropic effects driven by the same causal variant(s) (**Fig. 3C**). The observed *IFIH1* pleiotropic effects were further strengthened by the observation in the Genomics plc UK biobank data that the independent low-frequency *IFIH1* missense allele p.I923V (rs35667974-C, MAF=1.8%), previously reported to result in *IFIH1* LOF and to protect against T1D, vitiligo, psoriasis, and PsA, and to increase risk of UC, was also associated with increased risk of asthma (OR= 1.18, P = 1.1x10^-4^), appearing as the top asthma-associated SNP at the locus (**Fig. 3D** and **Fig. S5**). Together, these and previous findings establish *IFIH1* as a gene with an “allelic series”^36^ and further support the therapeutic hypothesis that inhibition of MDA5 may protect against autoimmune disease. However, our results also reveal the potential of MDA5 inhibitors to cause pulmonary ADEs and strengthen previous findings for an increased risk for colitis-related symptoms, endpoints that may limit the therapeutic window of MDA5 modulators and should be considered for monitoring in clinical trials.

**Fig. 3.**
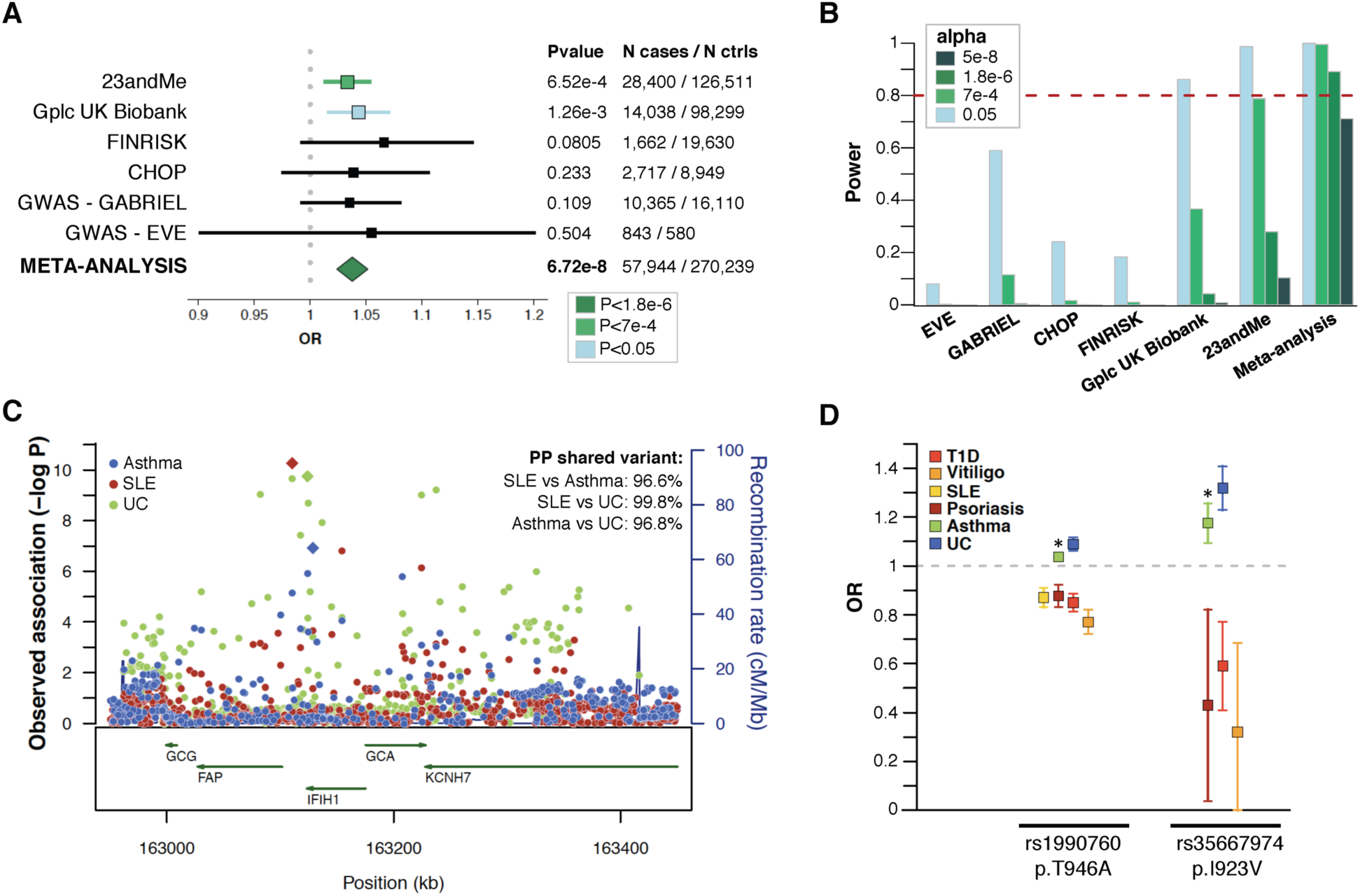
Pleiotropic effects of *IFIH1* LOF variants. (A) A significant association of *IFIH1* rs1990760-C (p.T946A) with increased risk of asthma was observed in the meta-analysis of PheWAS and GWAS results, with consistent effect estimate across the six cohorts tested. **(B)** Power estimation demonstrates the lack of power to detect an association at rs1990760-C in currently available asthma GWAS studies. Power to surpass various significance cutoffs (P<0.05; FDR<0.1, P<7e-4; study-wide significance after Bonferroni correction, P<1.8e-6; and genome-wide significance, P<5e-8) in the six cohorts was estimated using the frequency of the asthma risk allele (RAF=0.39), the odds ratio in the PheWAS/GWAS meta-analysis (OR=1.037), a disease prevalence of 8%, and the number of cases and controls in each of the cohorts. **(C)** Co-localization analysis demonstrates that the asthma, SLE and ulcerative colitis (UC) associations at the *IFIH1* locus are driven by a shared causal signal. PP, posterior probability. **(D)** Results from this study (*) combined with previously published findings suggest an allelic series of LOF *IFIH1* alleles decreasing the risk of various autoimmune diseases while increasing the risk of asthma and ulcerative colitis. Association results for the reported *IFIH1* loss-of-function alleles rs1990760-C (p.T946A) and rs35667974-C (p. I923V) are shown.

### Genetic efficacy and safety signals assist target prioritization for thromboembolism

Beyond informing on individual genes, we hypothesized that PheWAS might help prioritize targets among several candidates within a biological pathway. Factors XI, XII and plasma kininogen (encoded by *KNG1*) are members of the contact activation coagulation pathway^37^. Anticoagulation therapies directed against these factors are hypothesized to have improved therapeutic windows over current standard-of-care, which is accompanied by significant bleeding liabilities^38^. With the aim to estimate genetic risk-benefit profiles for the three candidate targets, we chose to interrogate three uncorrelated SNPs at the *F11*, *KNG1* and *F12* loci. These three SNPs had similar allele frequencies in Europeans, had previously been shown to impact FXI, FXII and/or KNG1 mRNA and/or protein levels, and are associated with activated partial thromboplastin time (aPTT), a biomarker of blood clotting, or venous thromboembolism (VTE), risk (**Supplementary Information** and **Table S1**). Carriers of the rs4253399-T allele, which reduces circulating FXI levels and increases aPTT, showed an expected lower risk for blood clots (OR=0.84, P=3.5x10-^25^), but no evidence for association with bleeding tendency (OR=1.04, P=0.35) (**Fig. 4**). In contrast, carriers of the *KNG1* allele rs5030062-A, which reduces plasma kininogen as well as circulating FXI, and inceases aPTT, showed both reduced blood clotting (OR=0.93, P=1.6x10^-4^) as well as increased bleeding liability (OR=1.14, P=4.1x10^-4^). A nominal association with both traits was found in carriers of the FXII levels-reducing and aPTT-increasing allele rs2731672-T (blood clots: OR=0.96, P=0.034; bleeding tendency: OR=1.09, P=0.039). By comparing these results with the effect of the three SNPs on aPTT (**Table S1**), our study suggests that, among the three factors tested, targeting FXI may yield the best compromise between thromboembolism risk reduction and increased bleeding liability, which is consistent with the outcomes of a recent phase 2 clinical trial^39^.

**Fig. 4.**
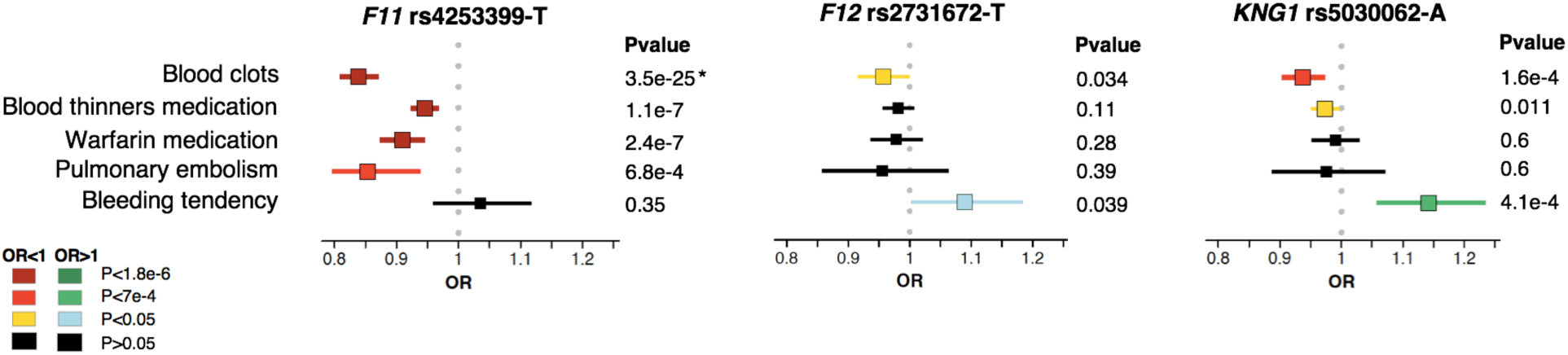
PheWAS association patterns of SNPs affecting genes in the contact activation coagulation pathway. Three SNPs known to affect plasma protein levels of FXI, FXII and KNG1, and previously reported as associated with partial thromboplastin time (aPTT) were interrogated in meta-PheWAS. Five phenotypes were observed as significantly associated (FDR<0.1) with at least one of the three SNPs: blood clots (known association with the *F11* SNP, *), blood thinners medication, warfarin medication, pulmonary embolism, and bleeding tendency. Effects of the aPTT-increasing alleles are shown.

## DISCUSSION

Our study investigates the utility of PheWAS for predicting therapeutic success of candidate drug targets nominated through human genetics. We focused on a selection of loci that GWAS have firmly established as associated with common immune-mediated, cardio-metabolic, or neurodegenerative human diseases, and where additional biological or genetic evidence supports candidate drug target genes within these loci as likely causing the disease associations. We analyzed SNPs impacting these targets for association with 1,683 disease endpoints captured in four large disease-agnostic “real-world” population cohorts that link genome-wide genotypes with various types of structured health information. Our PheWAS meta-analysis replicates 70% of the published GWAS associations at FDR<0.1, substantially surpassing performance of previous PheWAS in smaller cohorts^24^. Through joining PheWAS results with published GWAS data, we identified 10 novel SNP-phenotype associations that exceeded stringent significance thresholds for multiple test correction, as well as additional putative associations with therapeutically relevant clinical endpoints. For a subset of prominent early drug targets, our results support previous genetic evidence for efficacy in distinct common disease indications. Our analysis further proposes alternative indications as opportunities for drug repositioning, and predicts on-target adverse drug events that may warrant preclinical or clinical monitoring.

Among others, we discovered novel associations for p.I148M in *PNPLA3*. This is a common gain-of-function missense allele increasing the risk for a range of liver phenotypes, which suggested that pharmaceutical inhibition of PNPLA3 could be a viable strategy to treat or prevent liver diseases. While our PheWAS support this hypothesis and further expand the indication spectrum of a putative PNPLA3 inhibitor to T2D, they also uncovered associations with severe acne and high cholesterol, phenotypes that if observed only during a clinical trial might put a therapeutic program at risk.

We also identified a novel association of the *IFIH1* loss-of-function allele rs1990760-C (p.T946A) with risk of asthma. The rs1990760-C allele, which protects against autoimmune diseases and increases risk of ulcerative colitis, has been shown to decrease interferon (IFN) signaling and lower resistance to viral challenge^40^, while complete loss-of *IFIH1* function makes children susceptible to severe viral respiratory infections^41, 42^. The association of rs1990760-C with increased risk of asthma discovered in our meta-PheWAS is consistent with the observation that bronchial epithelial cells from asthmatics produce lower amounts of IFN-β during viral infections^43^, a finding that lead to inhaled IFN-β being tested in phase 2 clinical trials for the treatment of virus-induced asthma exacerbation^44^. Future studies will need to investigate the risk:benefit ratio of modulating MDA5 (encoded by *IFIH1)* for asthma relative to autoimmune disease.

While our study illustrates the power of systematically interrogating RWD cohorts to enrich target validation, it also emphasizes several opportunities to improve existing resources in order for PheWAS to become a routine tool in drug discovery and development. First, truly large, thoroughly phenotyped cohorts will be needed to adequately power PheWAS. Despite our study being conducted in more than 800,000 individuals, about one third of GWAS associations could not be replicated in the RWD cohorts at a stringent level of statistical significance due to an insufficient number of cases. In addition, PheWAS should considerably gain from improved ascertainment of phenotypes^45^. In our study, this is best reflected by an only modest replication rate, despite adequate power, for CD, UC and IBD endpoints that are closely related and difficult to discern in routine clinical settings. To better take these considerations and other characteristics of RWD cohorts (typical case:control ratio unbalance between phenotypes, and phenotype correlation) into account, novel statistical methods will be needed to better define significance thresholds and control type I error rates in PheWAS^46^. Second, our study highlights the challenge to systematically combine phenotypes from independent RWD cohorts. While we introduce the concept of “meta-PheWAS” and demonstrate that mapping phenotypes to interrogate independent PheWAS cohorts may considerably strengthen association signals, standardized terminology, automated phenotype extraction, and coordinated data management across healthcare institutions such as within the eMERGE network should help with better harmonization across cohorts in the future^9, 47^. A third challenge to the PheWAS approach is inherent to the current limitations of human genetics. Even when starting from a highly-annotated set of loci as in our study, PheWAS may lead to spurious associations that can only be ruled out through thorough follow-up^17^. We demonstrate this at the example of *LGALS3* and Parkinson’s disease. Access to genome-wide association results for systematic fine-mapping and co-localization analyses, functionalization of GWAS loci and the emergence of association data for intermediate phenotypes, e.g. at the protein level, will be needed to help narrow the gap between SNPs and candidate target genes in the future. Finally, a fourth challenge to broadly use PheWAS for drug development is to relate findings from germline variants that impact a target across an individual’s entire lifetime to success of an interventional trial with much shorter observation periods. In the end, many decisions to pursue or discontinue a therapeutic program may remain dependent on the specific risk/benefit ratio that quantitative genetics as applied here may help to predict, and the level of unmet clinical need.

Taken together, our study highlights PheWAS as a highly promising, yet largely untapped opportunity to use disease-agnostic “real world” cohorts for drug target validation. We provide several examples that illustrate PheWAS as a powerful strategy to help predict efficacy and unintended drug effects, which should ultimately help to develop better drugs. Whether PheWAS may truly impact decision-making during drug development will only become evident with either the emergence of ADEs in trials that genetics could have predicted, or reduced safety-related attrition rates for portfolios enriched in targets nominated through human genetics. The growing number of large-scale population cohorts that link genetic with extensive clinical data, together with an increased willingness across the borders of academia, biotech and the pharmaceutical industry to collaborate and share data, will provide opportunities to demonstrate that.

## MATERIALS AND METHODS

### SNP selection

In this study, we selected 25 SNPs that were significantly associated (P<5x10^-8^) in published GWAS with binary or quantitative phenotypes related to three main therapeutic areas: (auto)immune, cardiometabolic, or neurodegenerative diseases (**Supplementary Information**). These 25 SNPs had either been functionally validated in published studies, establishing the candidate target gene as causal for the risk of disease, or they were located within or near genes for which previous studies had generated convincing biological evidence to be of relevance for the respective clinical endpoint. The 25 SNPs were linked to 19 genes that were evaluated as candidate drug targets. Detailed information on the SNPs, candidate causal genes and their link to common human disease is provided in **Supplementary Information**. The list of SNPs and their known associated phenotypes is provided in **Table S1**.

### Study cohorts

We interrogated four large observational disease-agnostic RWD cohorts of subjects of European ancestry with genome-wide genotyped data linked to extensive phenotypic information (**Table 1**). All participants included in each of the four cohorts were unrelated individuals of European ancestry. Individual-level data from each cohort was analyzed independently, and the relevant summary statistics for the 25 SNPs were shared for further analysis. We restricted all cohorts to binary disease phenotypes with at least 20 cases per cohort.

#### 1) 23andMe

The 23andMe cohort comprised up to 671,151 participants and 654 binary disease endpoints derived from questionnaire-based self-reports^21^. Participants were restricted to a set of individuals who have >97% European ancestry, as determined through an analysis of local ancestry using a support vector machine (SVM) and a hidden Markov model (HMM) to assign individuals to one of 31 reference populations. A maximal set of unrelated individuals was chosen for each phenotype using a segmental identity-by-descent (IBD) estimation algorithm. Individuals were defined as related if they shared more than 700 cM IBD, including regions where the two individuals share either one or both genomic segments identical-by-descent. SNPs with Hardy-Weinberg equilibrium *P* < 10^−20^, call rate < 95%, or with large allele frequency discrepancies compared to European 1000 Genomes reference data were excluded. Participant genotype data were then imputed against the September 2013 release of 1000 Genomes Phase1 reference haplotypes^48^, using an internally developed phasing tool, Finch, which implements the Beagle haplotype graph-based phasing algorithm^49^, and Minimac2^50^.

#### 2) Genomics plc UK Biobank

The Genomics plc analysis of UK biobank cohort (referred to as ‘Genomics plc UK Biobank’) comprised 112,337 participants and 90 binary disease endpoints derived from questionnaire-based self-reports and medical interviews^10^. GWAS analyses were performed by Genomics plc using the interim data release (May 2015). QC followed the recommendations provided by UK Biobank. European ethnicity was defined as self-reported “white British” ethnic background, and confirmed by principal component analysis clustering. Samples with relatives (3rd degree or closer) were excluded. Imputation was carried out by the UK Biobank data providers using SHAPEIT3^51^, IMPUTE3^52^, and a reference panel combining the 1000 Genomes Phase 3^53^ and UK10K datasets^54^.

#### 3) FINRISK

FINRISK is a collection of cross-sectional population surveys carried out since 1972 to assess the risk factors of chronic diseases and health behavior in the working age population of Finland^22^. The FINRISK cohort comprised 21,371 Finnish participants and 269 binary disease endpoints derived from ICD codes grouping in Finnish national hospital registries and cause-of-death registry, and drug reimbursement and purchase registries. The FINRISK samples were genotyped using Illumina CoreExome, OMNIExpress, and 610K chips. After gender check, samples with genotype missing rate >5% or excess herozygosity (>4SD) were excluded. SNPs QC, including exclusion of SNPs with genotype missing rate >2%, minor allele frequency <1%, or Hardy-Weinberg equilibrium P value <1x10^-6^, was performed for each genotyping chip separately. Multidimensional scaling (MDS) components were estimated with PLINK v1.9^55^ from the LD-pruned genotype data where relatives with pi-hat>0.2 had been removed. Samples with non-Finnish ancestry observed as MDS outliers were removed. Imputation was performed with SHAPEIT^51^ and IMPUTE2^52^ using a reference panel combining information from the 1000 Genomes phase 3^53^ and 1,941 Finnish SiSu whole genome sequences^56^. Imputation was stratified based on genotyping chip.

#### 4) CHOP

The cohort from the Children’s hospital of Philadelphia (CHOP) comprised 12,044 pediatric patients and 870 binary disease endpoints derived from ICD9–CM codes using the ICD9-to-PheWAS codes mapping described by Denny *et al*^23, 57^. Subjects included in the CHOP PheWAS were genotyped on one of the following genotyping chips following the Illumina standard protocols: Illumina Human610-Quad version 1, Illumina 550K SNP array, or Illumina OmniExpress array. Samples with genotype call rate above 95% were included in the study. SNPs with genotype missing rate >5%, minor allele frequency <1%, and Hardy-Weinberg equilibrium P value <0.00001 were excluded. Principle component analysis (PCA) was performed using EIGENSTRAT^58^ on approximately 130,000 SNPs that had been pruned for linkage disequilibrium using PLINK v1.07^55^ and reference genotypes from the HapMap consortium^59^. Imputation was performed with SHAPEIT v2^51^ and IMPUTE2^52^ using the 1000 Genomes project phase 1 reference panel^48^. SNPs with with INFO scores <0.9 were excluded.

All the participants in the 23andMe, Genomics plc UK Biobank, FINRISK and CHOP cohorts provided written informed consent for participating in research studies. Blood samples were collected according to protocols approved by local institutional review boards. This research has been conducted using the UK Biobank resource under the Genomics plc project application number 9659.

In addition, with the aim to replicate novel associations identified in the RWD cohorts, we interrogated genome-wide summary statistics from 57 published GWAS, including 34 binary disease phenotypes, derived from a larger database that has been assembled and harmonized by Genomics plc (referred to as ‘Genomics plc GWAS’). The full list of studies in Genomics plc GWAS database and tested in this study is available in the **Supplementary Information**). These included checks to ensure consistency of the data, and alignment of alleles to the forward strand of the human reference sequence, with effects ascribed to the alternative allele. Effect size estimates for quantitative traits were rescaled relative to the residual variance. Summary-statistic imputation was applied to infer association evidence at common variants (minor allele frequency > 2%) in the 1000 Genomes EUR reference panel. Results for SNPs associated with the relevant phenotype with P < 0.05 were included in the meta-analysis.

Correlation between all GWAS was estimated to ensure that no GWAS included in the meta-analysis for a given phenotype presented overlapping samples. In addition, to further prevent GWAS results from overlapping samples to be meta-analyzed, only the most recent/largest study for a given disease was included in our analysis when several GWAS studies in the Genomics plc database investigated the same disease. Although we could not directly estimate potential overlapping samples between the different RWD cohorts, significant overlap is very unlikely based on the participants’ characteristics (**Table 1**).

### Identification of shared phenotypes

The phenotypic endpoints tested in the different RWD cohorts were derived from different sources (self-reports, medical interviews, WHO ICD codes, ICD9-CM codes) and named with different coding systems (e.g. clinical terms *versus* popular terms, abbreviations *versus* full names). In order to compare and combine results from the four RWD cohorts with published GWAS results from the Genomics plc database, we manually mapped the phenotypes. This step allowed us to identify 145 distinct phenotypes shared by at least 2 cohorts and with at least 20 cases in the independent cohorts (**Fig. 1**). The full list of mapped phenotypes is provided in **Table S2**. We note that, in each cohort some phenotypes were captured multiple times by different endpoints with slightly different definitions. In this case, only one endpoint per cohort was selected for meta-analysis.

### PheWAS and meta-analysis

Phenome-wide association analyses for each of the 25 SNPs were conducted in the 23andMe, Genomics plc UK biobank, FINRISK (PheWAS results release November 2016) and CHOP cohorts separately. Each SNP-phenotype association was tested independently (assuming an additive genetic model), using logistic regressions adjusted for age, gender, and principal components to adjust for population stratification. Genotyping batch and survey cohort were also included as covariates in the FINRISK PheWAS. We then performed two distinct analyses to 1) replicate known GWAS associations, and 2) to detect novel associations.

First, we meta-analyzed PheWAS results from the 4 RWD cohorts, to investigate the ability of these cohorts to replicate known GWAS associations. After harmonizing the effect alleles across the cohorts, fixed effect and random effect meta-analyses were performed using PLINK^55^. We then compared the meta-analysis association results with known significant SNP-phenotype associations from published GWAS, taking into account the statistical power to detect an association in the meta-analysis of the PheWAS results in the disease-agnostic RWD cohorts.

Second, we meta-analyzed results from the four disease-agnostic cohorts together with available GWAS results in order to detect novel associations. Meta-analysis was performed using PLINK as described above. Meta-analysis results at the 145 shared phenotypes were then combined with cohort-specific phenotype results from the 25 SNPs, resulting in 27,762 tests in total. False discovery rate (FDR) was calculated for the 27,762 tests combined using the Benjamini and Hochberg method using the R p.adjust method^60^. The threshold for significance in detecting putative novel associations was established using an FDR of 0.1, corresponding to P<7x10^-4^. Significance threshold based on Bonferroni correction was P=0.05/27,762 = 1.8x10^-6^. We note that Bonferroni correction ignores the correlation structure between the tested phenotypes or the fact that all the SNPs tested in this study are known to be associated with one or several phenotypes in published GWAS.

### Statistical power estimations

We estimated statistical power to detect an association with known associated phenotypes, based on the published effect size in the most recently published GWAS, the frequency of the associated SNP risk allele in the 1000Genomes EUR population, the number of cases and controls in the disease-agnostic cohorts, and assuming a phenotype prevalence of 1%^61^.

### Co-localization analyses

To distinguish true pleiotropic effects from multiple associations at the loci that are explained by different causal SNPs (and potentially incriminating different causal genes), we used association summary statistics available from published GWAS and applied a Bayesian test implemented in the R package ‘coloc’ to assess co-localization, i.e. the probability of sharing causal genetic variants between pairs of apparent pleiotropic phenotypes using association summary statistics at the loci of interest^27^. Co-localization analysis at the *LGALS3* locus was performed using meta-analyzed PD GWAS summary statistics from 23andMe published elsewhere (N cases=4,127, N controls=62,037)^25^, and galectin-3 plasma pQTL results in 3,562 blood donors^33^. Co-localization analysis at the *IFIH1* locus was performed using meta-analyzed SLE GWAS results from two independent published studies^62, 63^, meta-analyzed asthma GWAS summary statistics from 23andMe^64^ and the Genomics plc UK Biobank (unpublished), and published UC GWAS summary statistics^65^.

## Acknowledgements

We thank the research participants from the 23andMe, UK Biobank, FINRISK and CHOP cohorts for their contributions to this study. We would like to thank Jyoti Shah, Jennifer Pai, Mark Sharp, Hongjie Sun and Ian Wallace for their input on phenotype mapping.

## Author contributions

DD, HR, DFR, JM, JCM, AKC, AB, AD participated in the design and/or interpretation of the reported experiments or results. DD and HR drafted the manuscript. DD performed the phenotype harmonization, meta-analysis and follow-up analyses. RMP and CSF provided supervisory support. CT, the 23andMe Research team and DH participated in the acquisition and/or analysis of the 23andMe data. CSF, CCAS, CV, MEW and PD participated in the acquisition and/or analysis of the Genomics plc UK Biobank data. MA, HM, EK, MR, JH, NB, SJ, DGM, VS, SR, MJD, and AP participated in the acquisition and/or analysis of the FINRISK data. MM, PMAS and HH participated in the acquisition and/or analysis of the CHOP data.

